# Multiple drug resistance in hookworms infecting greyhound dogs in the USA

**DOI:** 10.1101/2021.04.24.438405

**Authors:** Pablo D. Jimenez Castro, Abhinaya Venkatesan, Elizabeth Redman, Rebecca Chen, Abigail Malatesta, Hannah Huff, Daniel A. Zuluaga Salazar, Russell Avramenko, John. S. Gilleard, Ray M. Kaplan

## Abstract

The hookworm *Ancylostoma caninum* is the most prevalent nematode parasite of dogs. Recently, we confirmed multiple-drug resistance (MDR) in several *A. caninum* isolates to all anthelmintic drug classes approved for the treatment of hookworms in dogs in the United States (USA). Cases of MDR hookworms appear to be highly overrepresented in greyhounds, suggesting that the MDR worms evolved on racing greyhound farms/kennels. The aims of this study were to evaluate the range of drug-resistant phenotypes and genotypes of the *A. caninum* infecting greyhounds. Fecal samples from recently retired greyhounds originating from geographically diverse areas of the USA were acquired from two greyhound adoption kennels, one active greyhound racing kennel, and three veterinary practices that work with adoption kennels. Fecal egg counts (FECs) were performed on fecal samples from 219 greyhounds, and despite almost all the dogs having been treated with one or more anthelmintics in the previous two to four weeks, the mean FEC was 822.4 eggs per gram (EPG). Resistance to benzimidazoles and macrocyclic lactones were measured using the egg hatch assay (EHA) and the larval development assay (LDA) respectively. We performed 23 EHA and 22 LDA on either individual or pooled feces, representing 81 animals. Mean and median IC_50_ and IC_95_ values for the EHA were 5.3 uM, 3.6 uM, and 24.5 uM, 23.4 uM respectively. For the LDA, mean and median IC_50_ values were 749.8 nM, >1000 nM respectively. These values range from 62 to 68 times higher than those we measured in our susceptible laboratory isolates. Pre-treatment fecal samples could not be obtained, however, post-treatment samples representing 219 greyhounds were collected. For samples collected <10 days post-treatment with albendazole, moxidectin, or a combination of febantel-pyrantel-moxidectin, the mean FEC were 349, 333, and 835 EPG, respectively. Samples collected 10-21 days post-treatment with albendazole, moxidectin, or pyrantel, yielded mean FEC of 1874, 335, and 600 EPG, respectively. Samples collected >21 days post-treatment with albendazole or moxidectin yielded mean FEC of 1819 and 1117 EPG, respectively. We obtained DNA from hookworm eggs isolated from 70 fecal samples, comprised of 60 individual dogs and 10 pools from multiple dogs. Deep sequencing of the isotype 1 β-tubulin gene revealed the presence of the F167Y (TTC>TAC) resistance polymorphism in 99% of these samples, with 69% having ≥75% resistant allele frequency. No resistance-associated polymorphisms were seen at any of the other β-tubulin codons previously reported as associated with benzimidazole resistance in Strongylid nematodes. These clinical, *in vitro*, and genetic data provide strong evidence that racing and recently retired greyhound dogs in the USA are infected with MDR *A. caninum* at very high levels in terms of both prevalence and infection intensity.

## 1. Introduction

The canine hookworm, *Ancylostoma caninum* is the most prevalent and important intestinal nematode parasite of dogs in the USA, with the prevalence depending on age, level of care and geographic location of the dog (Little et al., 2009). A recent study evaluating over 39 million fecal samples from 2012-2018, found that the prevalence of hookworms remained very stable from 2012-2014 at around 2%, but then from 2015 onwards, there was a steady yearly increase, with an overall increase of 47% by 2018 (Drake and Carey, 2019). Moreover, in a study assessing intestinal parasites from 3006 dog fecal samples collected in 288 off-leash dog parks across the USA in 2019, the prevalence of *A. caninum* was 7.1% (Stafford et al., 2020). Interestingly, this prevalence is more than twice as high as that reported for 2018 by (Drake and Carey, 2019), and is more than 70% higher than the mean prevalence for 2017-2019 reported by (Sweet et al., 2021). Taken together these data suggest that hookworm prevalence is rapidly increasing, and that dogs that visit dog parks are at a higher risk of infection.

Anthelmintic drugs currently approved for the treatment of *A. caninum* in the United States include, febantel and fenbendazole, moxidectin and milbemycin oxime, and pyrantel, of the benzimidazole, avermectin/milbemycin, and tetrahydropyrimidine classes, respectively. In registration studies, febantel, moxidectin and milbemycin oxime all demonstrated efficacies of >99% (F.D.A, 1994, 1998, 2006), fenbendazole demonstrated efficacy of >98% (F.D.A, 1983) and pyrantel demonstrated a slightly variable efficacy, with a mean across studies of approximately 94%, where more than half of those studies yielded >99% (F.D.A, 1993).

Hookworms are blood-feeding nematodes that use a cutting apparatus to attach to the intestinal mucosa and submucosa, and contract their muscular esophagus to create negative pressure, which sucks a plug of tissue into their buccal capsules (Hotez et al., 2004). Bleeding is facilitated by both mechanical damage and chemical action by hydrolytic enzymes that cause rupture of capillaries and arterioles (Stassens et al., 1996). Pathological consequences of infection in dogs include iron-deficiency anaemia, hypoalbuminemia, and an enteritis characterized by diarrhoea, that may contain fresh (haematochezia) or digested blood (melena) (Epe, 2009; Kalkofen, 1987; Taylor et al., 2016).

In the past few years, there is empirical evidence that veterinarians are diagnosing increasing numbers of cases of persistent hookworm infections, primarily in greyhound dogs, that appear refractory to typical anthelmintic therapy. Recent work in our laboratory confirmed that many, if not most, of these persistent hookworm cases are likely due to multiple-drug resistance (MDR) in *A. caninum*, with retired racing greyhounds highly over-represented among the cases reported to our laboratory. Our laboratory established one of these *A. caninum* isolates (Worthy), which we obtained from a recently adopted retired racing greyhound dog. In a controlled efficacy study, we confirmed high levels of resistance to all classes of drugs approved for treatment of hookworm in dogs; fenbendazole, pyrantel pamoate and milbemycin oxime yielded efficacies of 26%, 23% and 9%, respectively (Jimenez Castro et al., 2020). *A. caninum* is also zoonotic in humans, and MDR *A. caninum* will not respond to usual anthelmintic treatments administered by physicians.

Greyhound racing was once very trendy and profitable in the USA. At the sport’s peak in 1991, dog racing was rated the sixth most popular sporting activity in the USA, was legal in 19 states, and generated around 100,000 jobs with wager revenues of $3.5 billion USD (Theil, 2021). At that time there were 38,000 individual pups and 5,700 registered racers, but by 2020 those numbers had dramatically decreased to 4,300 and 850, respectively (Gartland, 2021). Kansas, the state with the most greyhound breeding farms, had 274 greyhound breeding farms in the 1990’s, but by 2015 this number had decreased by more than half to 130 (Hall, 2016), and continues to fall. These farms tend to have large dog populations; more than 60% and 80% of all racing greyhounds come from farms with >250 dogs and >100 dogs, respectively (Hall, 2016).

In November, 2018, voters in FL passed Amendment 13, a constitutional amendment which banned wagering on live dog races, including greyhound racing in the state as of January 1st, 2021 (State, 2018). In 2018, 65% of the greyhound tracks in the USA were in Florida. However, with the closure of these tracks and others in several other states in the past few years, to our knowledge there currently are only 7 tracks in 5 states remaining. This change likely represents the beginning of the end for greyhound racing in the USA. In parallel, a large network of greyhound adoption groups has been active for many years, having over 160 organizations across the USA and Canada with the majority being in FL followed by NY and OH. This results in thousands of retired racing greyhounds being adopted as pets every year (Lord et al., 2007), and with the demise of the greyhound racing industry, the number of adoptions are rapidly increasing. Thus, it is important for the health of both racing greyhounds and pet dogs to determine the extent of the MDR hookworm problem in racing greyhounds.

The aims of this study were to investigate the prevalence of infection, the range of *in vitro* and *in vivo* drug susceptible/resistant phenotypes, and the frequency of benzimidazole-resistant beta-tubulin genotypes in greyhound dogs infected with *A. caninum*.

## 2. Materials and Methods

### 2.1 Sample collection

From February 2019 to February 2020, fecal samples were acquired from two greyhound adoption kennels located in Birmingham, AL, and Dallas, TX, one active greyhound racing kennel in Sanford, FL, and three veterinary practices located in Acworth, GA, Columbia, SC, and St. Petersburg, FL USA that work with greyhound adoption organizations. Most samples were collected from individual dogs, but from the Sanford, FL site, only anonymous samples from the ground were available. The dogs residing in these kennels originated from 16 different locations in 8 different states. These included five breeding farms located in KS, CO, AR, TX, or OK, and 11 racing tracks located in AL, FL, AR, or WV.

### 2.2 *In vitro* assays

To evaluate drug response phenotypes, the egg hatch assay (EHA) and larval development assay (LDA) were used for benzimidazoles (BZs), and macrocyclic lactones (MLs), respectively as previously described (Jimenez Castro et al., 2019). The concentration ranges for ivermectin aglycone (1.9 – 1000 nM) in the LDA, and thiabendazole (0.075 – 40 µM) in the EHA were selected based on our previous work to permit the discrimination of susceptible vs resistant isolates of *A. caninum* (Jimenez Castro et al., 2019). Eggs were isolated using 50 ml tubes containing charcoal granules (0.5 – 1 cm) and specialized lids containing a filter that could attach to 15 ml centrifuge tubes. Five to ten grams of feces and approximately 15 ml of water were added to the 50 ml tube containing approximately 5 grams of charcoal and vigorously shaken to break up the feces. The lid of the 50 ml tube was removed and replaced with the specialized lid, and a 15 ml tube was attached to the other end. The apparatus was then shaken again which allowed the filtered fecal suspension to fill the 15 ml tube, which was then centrifuged at 240 x *g* for 10 mins. The supernatant was discarded, 10 ml of sodium nitrate (Feca-Med®, Vedco, Inc. St. Joseph; MO, USA specific gravity = 1.2) were added and the tube was vortexed to disperse the fecal material. The tube was then centrifuged again at 240 x *g* for 10 mins. Following centrifugation, the supernatant containing the eggs was passed through a 20 µm sieve, rinsed with distilled water, transferred to a new tube, and then the volume was adjusted to yield 50-60 eggs per20 μl using distilled water. If insufficient eggs were recovered to perform both the EHA and the LDA, then only the EHA was performed.

### 2.3 *In vivo* measurements

Every dog sampled in this study was treated with anthelmintics at regular intervals; therefore, all of the samples were collected relatively recently post-treatment. Samples were refrigerated immediately after collection and shipped to the Kaplan lab at the University of Georgia in a container with ice packs. In order to account for the differences in timeframe since the previous anthelmintic treatment, dogs were assigned to one of three categories based on the following biological factors: (A) = <10 days, as this can be too soon to measure an accurate fecal egg count reduction (FECR) and can lead to false positives due to temporary inhibition of egg production (Jimenez Castro et al., 2019; Jimenez Castro et al., 2020), (B) = 10 – 21 days, as this would be an optimal timeframe for measuring the FECR, and (C) = > 21 days, as there is the possibility that eggs could be shed from reactivated encysted/arrested larvae that migrated to the small intestine and completed development to sexually mature adults following the anthelmintic treatment (due to “larval leak”) (Jimenez Castro and Kaplan, 2020). Fecal egg counts (FEC) were performed using the Mini-FLOTAC (University of Naples Federico II, Naples, Italy) procedure with a detection threshold of 5 EPG (Lima et al., 2015; Maurelli et al., 2014), adding two grams of feces to 18 ml of sodium nitrate (Feca-Med®, Vedco, Inc. St. Joseph; MO, USA specific gravity = 1.2). Positive samples were defined as having a FEC of ≥ 5 EPG. All anthelmintic treatments were administered by either kennel or veterinary practice personnel. Where products approved for use in dogs were used, treatments were administered according to label instructions; these included febantel-pyrantel pamoate, Drontal® Plus (Elanco, Greenfield, IN), moxidectin, Advantage® Multi (Elanco, Greenfield, IN) and pyrantel pamoate, Nemex-2® (Zoetis, Kalamazoo, MI). In some instances, products labelled for large animals were used; these included moxidectin, Quest® Plus (Zoetis, Kalamazoo, MI) and albendazole, Valbazen® (Zoetis, Kalamazoo, MI). These products were administered orally at 3.3 mg/kg and 19 mg/kg, respectively. In some cases, compounded drugs were used, such as the PPM Triwormer (Roadrunner pharmacy, Phoenix, AZ).

### 2.4 *Ancylostoma caninum* isotype□1 beta□tubulin deep amplicon sequencing

#### 2.4.1 DNA preparation

After setting up the *in vitro* assays, the remaining eggs were transferred to 2 ml cryotubes (Sigma-Aldrich, St. Louis, MO), suspended in a final concentration of 70% ETOH and stored at -80°C until further use. DNA lysates were prepared from individual or pooled egg samples. Briefly, 3 freeze-thaw cycles were carried out at -80°C and at 55°C respectively, followed by adding 180 μL of DirectPCR (Cell) Lysis Buffer (Catalog No. 301-C, Viagen Biotech, St. Louis, MO) and 20 μL of Proteinase K (Catalog No. 19133, QIAGEN, Hilden, Germany). Samples were then incubated for at least 12h at 65°C, then 1h at 95°C, and were then cooled to 4°C. DNA was purified from the crude DNA lysates using the QIAGEN QIAmp DNA mini kit (Cat# 51306), following the manufacturer’s recommended protocol, and stored at -80°C.

#### 2.4.2 Deep-amplicon sequencing assay analysis

Deep amplicon sequencing assays developed to evaluate the frequency of non-synonymous single nucleotide polymorphisms (SNP) at codons 167, 198 and 200 of the *A. caninum* isotype-1 β -tubulin gene were applied to 70 samples ranging from 200 - 20,000 eggs with an average of 978 eggs from the two adoption kennels, one active racing kennel, and from one of the veterinary practices. Using adapted primers suitable for Illumina next-generation sequencing, two separate regions of the *A. caninum* isotype-1 β -tubulin gene, comprising 293 bp and 340 bp which encompass codon 167, and codons 198 and 200, respectively, were PCR amplified (Jimenez Castro et al., 2019). The following PCR conditions were used: 5µL KAPA HiFi Hotstart Fidelity Buffer (5X) (KAPA Biosystems, USA), 1.25µL Forward Primer (10µM), 1.25µL Reverse Primer (10µM), 0.75µL dNTPs (10µM), 0.5µL KAPA HiFi Polymerase (0.5U), 0.1µL Bovine Serum Albumin (Thermo Fisher Scientific), 14.15µL H2O, and 2µL of DNA lysate. The thermocycling parameters were 95 □ for 3 min, followed by 45 cycles of 98 □ for 20 s, 65 □ for 15 s, and 72 □ for 30 s, followed by 72 □ for 2 min. Sample purification and addition of barcoded primers followed the protocols defined in (Avramenko et al., 2019). Library preparation was as previously described and library sequencing performed using the Illumina MiSeq platform with the 2 × 300v3 Reagent Kit (Illumina Inc., San Diego, CA, USA) (Avramenko et al., 2015). For the fragment encompassing codon 167, two independent PCR reactions were performed on all 70 samples and the libraries were sequenced in two independent sequencing runs using the Illumina MiSeq platform with the 2 x 300 v3 Reagent Kit.

#### 2.4.3 Sequence analysis

Cutadapt v3.2 (Martin, 2011) was used to remove the *A. caninum* forward and reverse primer sequences. Following adapter trimming, all the forward and reverse reads were processed using the DADA2 bioinformatic pipeline to obtain Amplicon Sequence Variants (ASVs) (Callahan et al., 2016). During the quality filtering step of the DADA2 pipeline, the default setting with the following additional settings were used: (i) forward and reverse reads were trimmed to a length of 280 bp and 190 bp, respectively and (ii) reads shorter than 50 bp or with an expected error of > 1 or > 2 in the forward and reverse reads respectively were removed. The DADA2 algorithm was then applied to the filtered and trimmed reads to identify the ASVs. Following this, the overlapping forward and reverse reads were merged, allowing a maximum mismatch of 4 bp in the overlap region. The ASVs generated using the DADA2 pipeline were then aligned to the *A. caninum* isotype-1 β -tubulin reference sequence (Genbank Accession: DQ459314.1) using a global (Needleman-Wunsch) pairwise alignment algorithm without end gap penalties. Following alignment, the ASVs were discarded if they were <180 bp or >350 bp long, or if they had a percentage identity <70% to the reference sequence, or if the ASVs had fewer than 200 reads in a sample, or if they were not present in two or more samples. This additional filtering ensures the removal of spurious sequences. The codons 167, 198 and 200 were then analyzed for the presence of any variants resulting in non-synonymous changes. The MUSCLE alignment tool was used to align the filtered ASVs from both the fragments with isotype-1 and 2 β -tubulins from other nematodes present in Clade V of the nematode phylogeny. Using the Geneious tree builder, a neighbour-joining tree, utilizing the Jukes Cantor tree building method, was constructed from the trimmed alignment, and having *H. contortus* isotype-3 β -tubulin as the outgroup (Genbank Accession: HE604101) and 2000 bootstrap replicates.

### 2.5 Data analyses

Dogs treated solely with albendazole and moxidectin were represented in each of the post-treatment timeframe categories, therefore for each drug individually, statistical analyses were performed to determine if the FEC of the dogs differed between those categories. For this a Kruswal-Wallis test was performed for the overall comparison with a Benjamini-Krieger-Yekutielli procedure for individual pairwise comparisons. For the EHA and LDA, dose-response analyses were performed after log transformation of the drug concentrations and constraining the bottom value to zero. The top parameter for samples that did not reach 100% inhibition was constrained to 100 to avoid introducing artificial bias to the model. Data were then fitted to a four-parameter non-linear regression algorithm with variable slope. The IC_50_ or IC_95_ values, which represent the concentration of drug required to inhibit hatching (EHA) or development to the third larval stage (LDA) by 50% or 95% of the maximal response were calculated.

The software https://epitools.ausvet.com.au/ was used to calculate the prevalence along with 95% confidence intervals. For comparing the infection prevalence between states, a Chi-square test was performed for the overall comparison, followed by a Fisher’s exact test with a Bonferroni procedure for individual pairwise comparisons between states. All analyses were designed to maintain the overall type I error rate at 5%. To quantify the relationship between the EHA and the deep-amplicon sequencing assay to measure levels of BZ resistance, a Spearman correlation analysis was performed comparing both IC_50_ and IC_95_ values with the F167Y SNP frequencies. All statistical analyses were performed in GraphPad Prism® version 9.0.2, GraphPad Software, San Diego, CA, USA.

## 3. Results

### 3.1 Fecal egg count data

171 of the 219 fecal samples from racing or recently retired greyhounds were positive for hookworm eggs, yielding an overall prevalence of 79%, with a mean FEC of 822.4 EPG (Table 1). The prevalence of infection for the Birmingham, AL (*p* = 0.0063) and Sanford, FL (*p* < 0.0001) kennels were significantly higher than for the Dallas, TX kennel. Percent reductions in FEC following treatments could not be calculated since pre-treatment FEC were not available. However, in the 125 (57.3%) samples collected 2 – 21 days post-treatment, the mean FEC was 721 EPG, indicating a major lack of efficacy across the different treatments. For the group of samples collected following moxidectin treatment, samples collected > 21 days post-treatment had a statistically significant higher mean FEC when compared to the other two categories. For the group of samples collected following albendazole treatment, the 10-21 and >21 days categories had a statistically significant higher mean FEC when compared to the <10 days. Interestingly, a single group of samples from nine dogs recently acquired from a breeding farm in KS had <5 EPG following treatment. For all other sites, mean FEC were 330 EPG or greater (Table 2).

**Table 1.** Mean fecal egg count (FEC) data and percent prevalence of hookworm infections from 219 greyhound dog samples obtained from two greyhound adoption kennels, one greyhound racing kennel, and veterinary practices. Data from the three veterinary practices were combined for reporting purposes. Positive samples were defined as having a FEC of > 5 eggs per gram (EPG). The software https://epitools.ausvet.com.au/ was used to calculate the prevalence along with 95% confidence intervals.

| **Adoption kennel** | **FEC (EPG)** | **Prevalence (%) (95%CI)\*** | **Total # of samples** |
| --- | --- | --- | --- |
| **Birmingham, AL** | 527.8 | 76.3 (66.2,84.7)<sup>a</sup> | 80 |
| **Sanford, FL** | 1288.6 | 87.5 (79.1,93.5)<sup>a</sup> | 80 |
| **Dallas, TX** | 683 | 48.1 (30.1,66.5)<sup>b</sup> | 27 |
| **Veterinary practices<sup>1</sup>** | 529.3 | 90.6 (77.5,97.6) | 32 |
| **Overall means** | 822.4 | 79 (73.3,84.0) | 219 |

<sup>1</sup> Three veterinary practices located in Acworth, GA, Columbia, SC, and St. Petersburg, FL
\*Shared superscripts denote non-significant differences

**Table 2.** Mean fecal egg counts (FEC) in eggs per gram (EPG) with the standard error of the mean (SEM) of fecal samples from greyhound dogs. Samples were obtained at varying intervals following treatments with several different anthelmintics. State of origin of the dogs is provided, and where more than one city was represented and city was known, a subscript letter indicates the number of cities represented. Timeframes since the previous anthelmintic treatment were categorized as: (A): <10 days, (B): 10 – 21 days, and (C): > 21 days. Dogs treated solely with albendazole and moxidectin were represented in each of the post-treatment timeframe categories, therefore for each drug individually, statistical analyses were performed to determine if the FEC of the dogs differed between those categories.

| Treatment | No. of greyhounds | Categories post-treatment* | FEC (SEM) (EPG) | State of origin<sup>5</sup> |
| --- | --- | --- | --- | --- |
| **Single drugs** |  |  |  |  |
| Moxidectin<sup>1</sup> | 20 | A<sup>a</sup> | 333.3 (108.9) | AL, FL<sub>1</sub>, FL<sub>2</sub>, FL<sub>3</sub>, AR, TX, WV, KS |
| Moxidectin<sup>1</sup> | 41 | B<sup>a</sup> | 334.6 (93.5) | AL, AR, FL<sub>1</sub>, FL<sub>2</sub>, FL<sub>4</sub>, WV, AR, CO, TX |
| Moxidectin<sup>1</sup> | 53 | C<sup>b</sup> | 1117 (189.4) | AL, AR, FL<sub>1</sub>, FL<sub>3</sub>, FL<sub>4</sub>, FL<sub>5</sub>, FL<sub>6</sub>, WV |
| Pyrantel pamoate<sup>2</sup> | 3 | C | 600 (263.2) | FL, AL |
| Albendazole<sup>3</sup> | 20 | A<sup>c</sup> | 349.3 (150) | AR, FL |
| Albendazole<sup>3</sup> | 21 | B<sup>d</sup> | 1874 (396.4) | AR, FL |
| Albendazole<sup>3</sup> | 10 | C<sup>d</sup> | 1819 (524.1) | FL |
| **Combinations** |  |  |  |  |
| Febantel-pyrantel-moxidectin<sup>4</sup> | 20 | A | 834.8 (271.9) | AL, AR, AR<sub>1</sub>, FL<sub>1</sub>, FL<sub>3</sub>, FL<sub>4</sub>, FL<sub>7</sub> |
| Moxidectin<sup>1</sup> + pyrantel<sup>2</sup> | 9 | C | 0 | KS |

<sup>1</sup> Quest® Plus
<sup>2</sup> Nemex®-2
<sup>3</sup> Valbazen®
<sup>4</sup> Drontal® Plus and Advantage® Multi
<sup>5</sup> FL<sub>1</sub> = Palm Beach, FL<sub>2</sub> = Daytona Beach, FL<sub>3</sub> = Jacksonville, FL<sub>4</sub> = Naples-Ft. Myers, FL<sub>5</sub> = Sanford, FL<sub>6</sub> = Clermont, FL<sub>6</sub> = Pensacola, AR<sub>1</sub> = West Memphis
\*For moxidectin and albendazole, shared superscripts denote non-significant differences

### 3.2 *In vitro* assays

EHA and LDA were performed on 35 samples, yielding dose-response data on 23 and 22 samples, respectively, which represented samples collected from 81 greyhounds (Fig. 1).

**Fig. 1.**
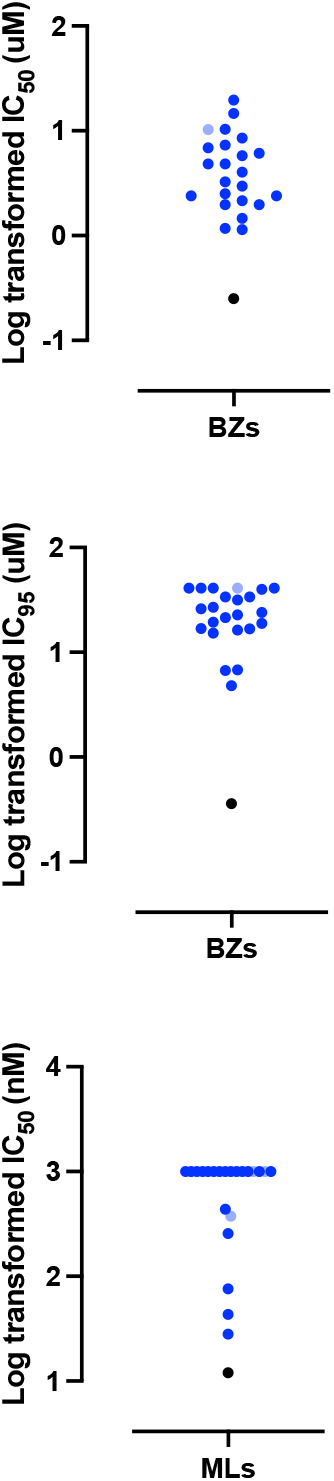
Scatterplots of greyhound samples showing the log transformed Egg Hatch Assay (EHA) IC_50_ (a) and IC_95_ (b) values, and the Larval Development Assay (LDA) IC_50_ (c) values for the benzimidazoles (BZs) and macrocyclic lactones (MLs), respectively. Each dark blue and light blue dot represent an assay performed on an individual or a pooled sample, respectively. The black dot represents the value of our susceptible laboratory isolate for reference. Dose-responses were analysed using the variable slope nonlinear regression model analysis contained in GraphPad 9.0.2.

Mean and median IC_50_ and IC_95_ values were 5.3 uM, 3.6 uM, and 24.5 uM, 23.4 uM for the EHA, respectively. For the LDA, mean and median IC_50_ values were 749.8 nM, >1000 nM, respectively (Table 3). IC_95_ values were calculated for the LDA (Table 3), but since the IC_50_ were greater than the highest concentration tested in the majority of samples, the mean and median values for IC_95_ have no real usefulness and are not reported.

**Table 3.**
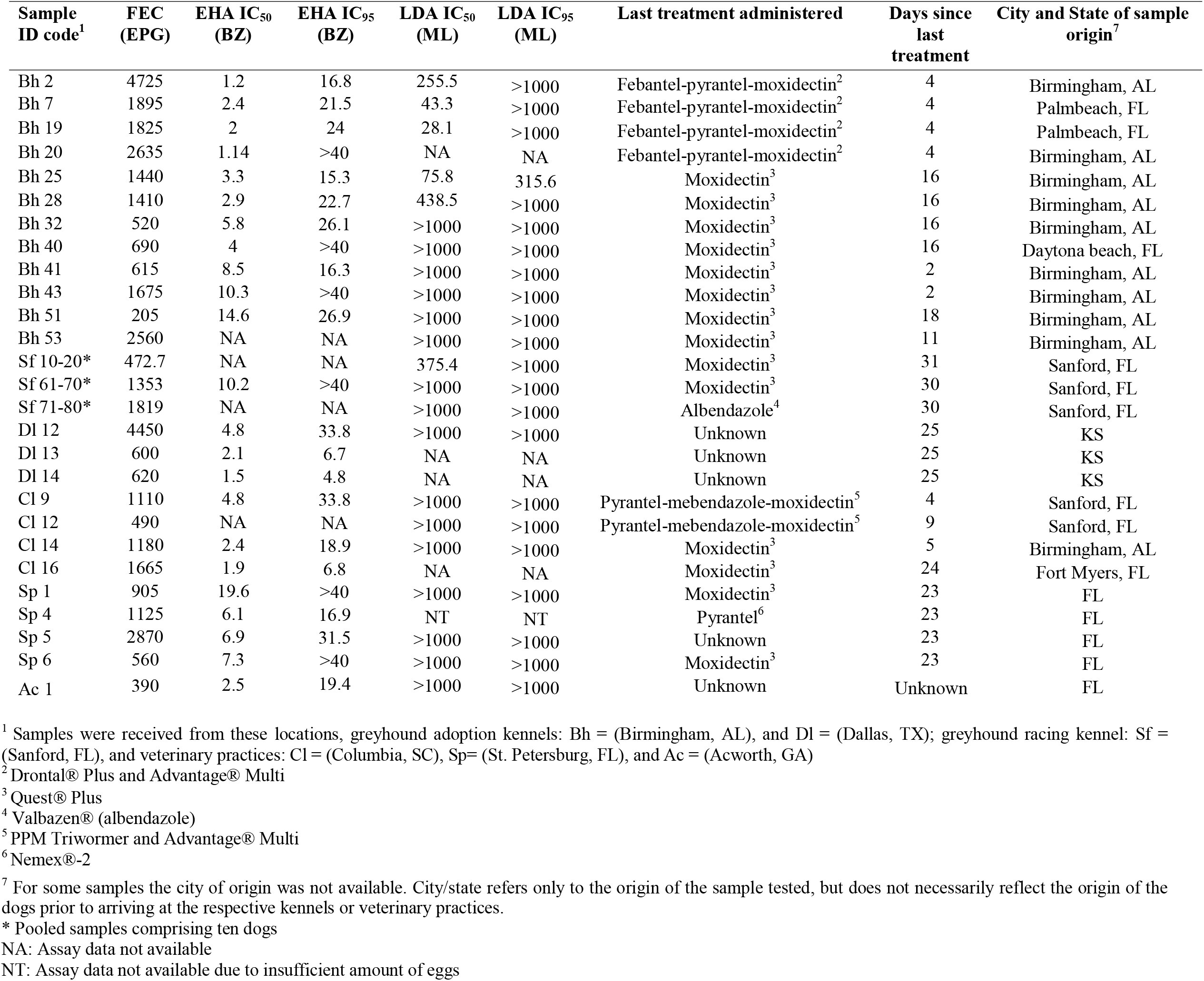
Geographic origin of the dogs, fecal egg counts (FEC) in eggs per gram (EPG), egg hatch assay (EHA) IC_50_ and IC_95_ values (uM), larval development assay (LDA) IC_50_ values and IC_95_ values (nM), last anthelmintic treatment administered, and days from treatment to sample collection. All dose-response analyses were performed after log transformation of the drug concentrations and constraining the bottom value to zero. Data were then fitted to a four-parameter non-linear regression algorithm with variable slope.

### 3.3 Relative frequencies of the isotype-1 β-tubulin benzimidazole resistance associated polymorphisms

The three codons in the isotype-1 β -tubulin gene known to have BZ resistance-associated polymorphisms (167, 198 and 200) in Strongylid nematodes were examined using deep-amplicon sequencing. The average read depth for the fragment containing codon 167 was ∼10,400, ranging between 3,180 to 24,732 reads across the samples. For the fragment containing codons 198 and 200, the average read depth was ∼22,700, ranging between 7,890 to 37,060 reads. Only the F167Y (TTC>TAC) resistance polymorphism was detected and this was present in 99% (69/70) of the samples that were sequenced, and this polymorphism was found at high frequencies in most positive samples (Fig. 2). In 48 out of the 70 samples (69%), the frequency of the resistant allele was ≥75%. In 29% of the samples, and in at least one sample from each greyhound kennel, the allele frequency was ≥ 90%. All greyhound kennels had samples with at least a 60% frequency of the F167Y SNP. Only 7% of the samples had <25% of the resistant allele, and only one sample had 100% frequency of the susceptible allele. These frequencies were consistent between the two independent sequencing runs for the fragment containing codon 167. The ASVs for the amplicons spanning codons 167 and codons 198 and 200 were aligned to other nematode β-tubulins using MUSCLE alignment tool. This alignment was trimmed, and a neighbour-joining tree was constructed using Geneious tree builder with *H. contortus* isotype-3 β-tubulin (Genbank accession: HE604101). All the ASVs formed a monophyletic cluster with *Ancylostoma* isotype-1 β-tubulin (supplementary figure 1 and 2).

**Fig. 2.**
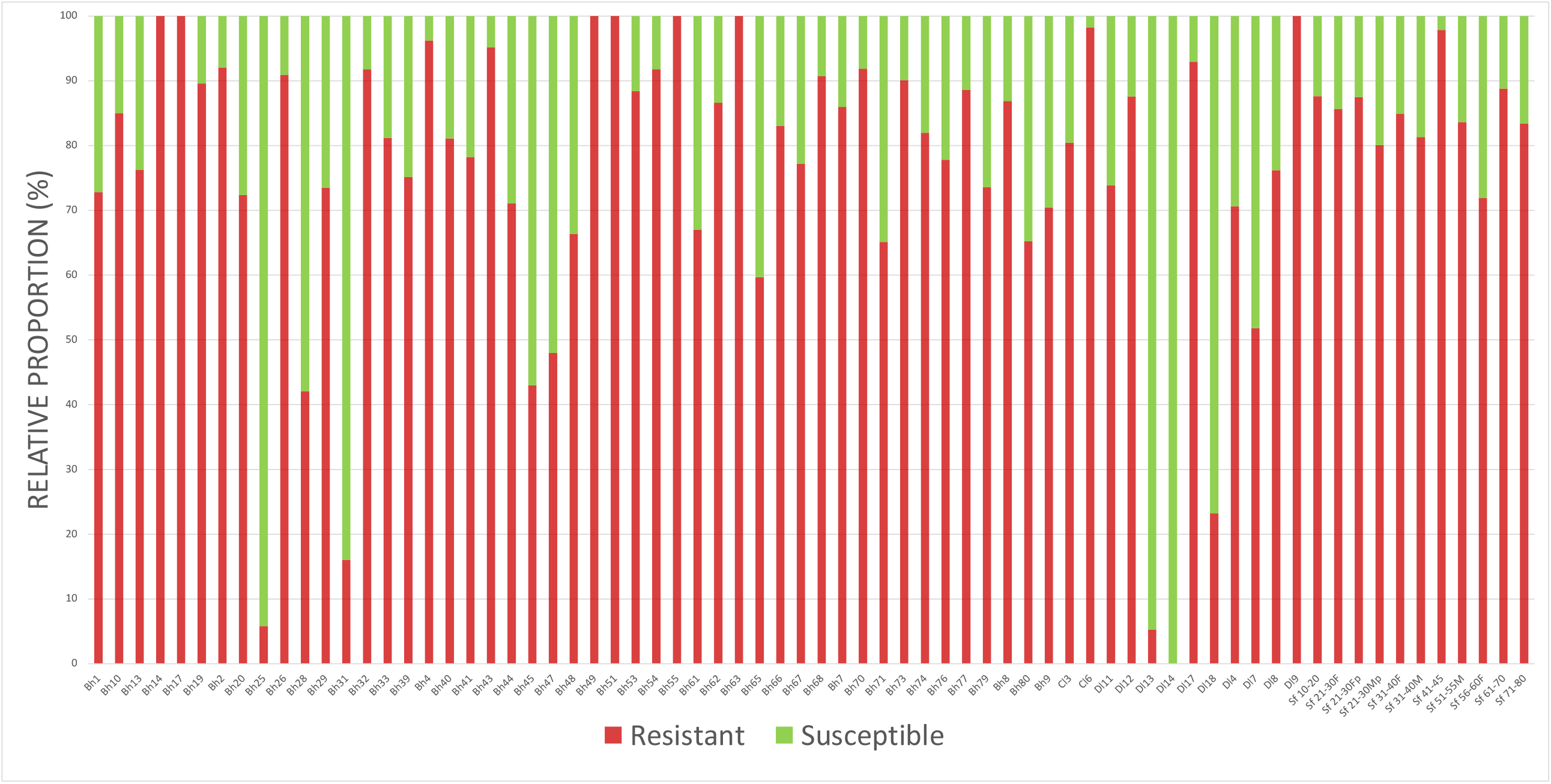
The relative proportions of isotype-1 beta-tubulin alleles encoding resistance conferring polymorphisms at F167Y vs wild type susceptible as measured by deep-amplicon sequencing in 70 samples from greyhounds that originated from 16 different locations in 8 different states.

### 3.4 Comparison of F167Y (TTC>TAC) frequency and EHA phenotype

Both EHA (phenotypic) and β-tubulin allele (genotypic) data were only available for 15 samples. There was not a statistically significant correlation between the IC_50_ and resistant SNP F167Y allele frequency (*p* = 0.08), however, there was a significant correlation with the IC_95_ (*p* = 0.04) (Fig. 4).

**Fig. 3.**
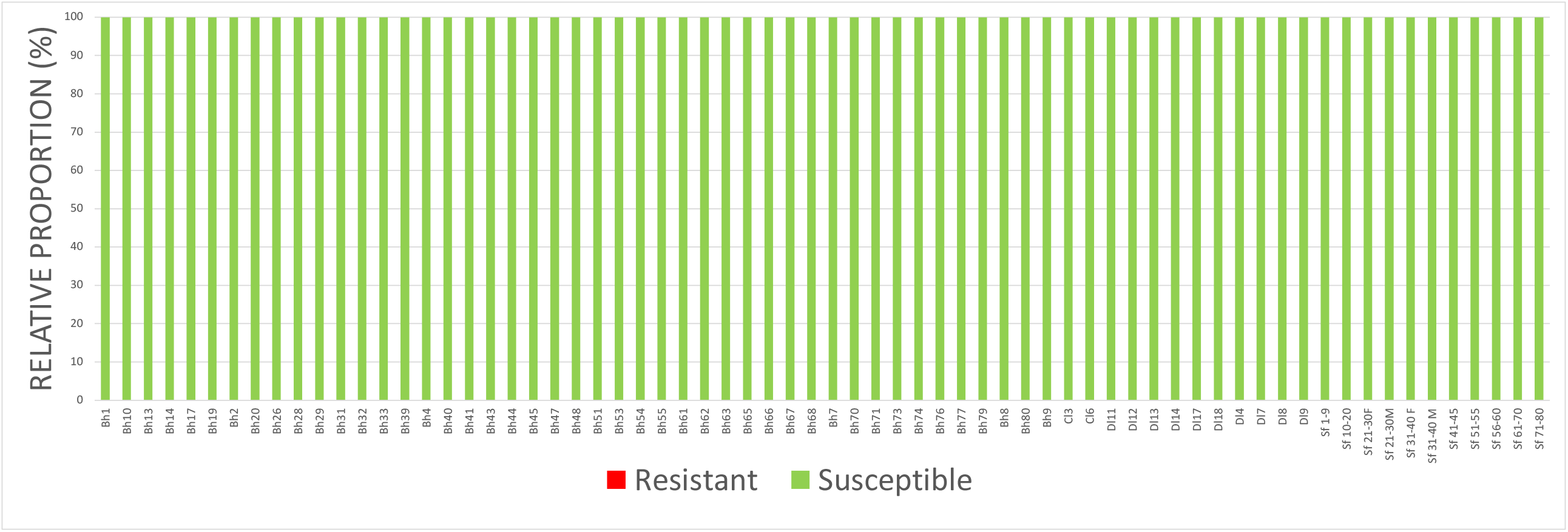
The relative proportions of isotype-1 beta-tubulin alleles in the fragment containing codons 198 and 200 as measured by deep-amplicon sequencing in 70 samples from greyhounds that originated from 16 different locations in 8 different states.

**Fig 4.**
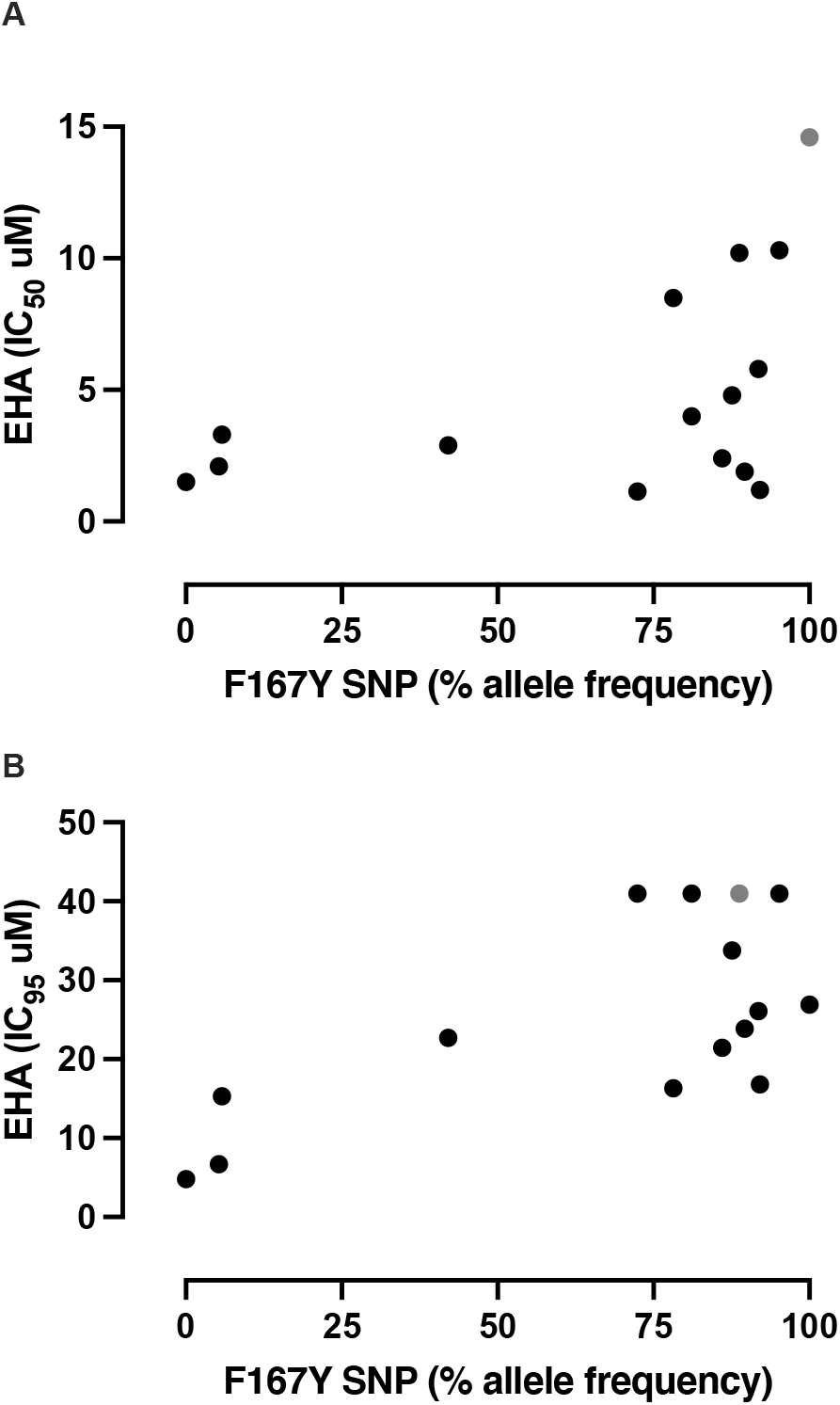
Scatterplots of EHA IC_50_ values (A) or IC_95_ values (B) vs. F167Y SNP frequency based on deep-amplicon sequencing. Only 15 samples had results from both assays. The gray dot represents a pooled sample. The highest concentration tested in the EHA was 40 uM, therefore the four values with IC_95_ of 40 uM likely would have been greater if higher concentrations were tested.

## 4. Discussion and conclusions

The present study provides strong and conclusive evidence that racing greyhounds in the USA are infected with MDR *A. caninum* at a very high prevalence, and with wide geographic distribution. Very high IC_50_ and IC_95_ values were measured for both the benzimidazoles and macrocyclic lactones when compared to the susceptible isolate from our previous work (Jimenez Castro et al., 2019), indicating that almost every sample was resistant to both drugs. The F167Y (TTC>TAC) benzimidazole resistance polymorphism was detected, in 99% of the samples, and at high frequencies in more than 2/3 of the samples. All three greyhound kennels had samples with at least a 60% frequency of the F167Y SNP, and every sample from the Sanford, FL site had at least a 70% frequency of the resistant SNP. These data are consistent with levels we reported for our MDR (Worthy) lab isolate, which yielded F167Y SNP frequencies of 87.6 - 94.5% over several different passages (Jimenez Castro et al., 2019). Furthermore, in a controlled efficacy study using Worthy 4.1F3P, we measured an efficacy of 26% for fenbendazole, confirming that this high F167Y SNP frequency was associated with a very low *in vivo* efficacy (Jimenez Castro et al., 2020).

Although additional genetic analyses are required for confirmation, the available clinical and genetic evidence strongly suggests that these MDR *A. caninum* evolved on greyhound breeding farms and kennels. Thus, it is germane to this issue to examine the clinical and epidemiological factors that may be responsible for the development of these MDR worms, and to hypothesize why this problem became so severe before it was recognized. *Ancylostoma caninum* is the most prevalent parasitic nematode in racing greyhounds (Ash et al., 2019; Jacobs and Prole, 1976), and this is attributed to the near constant exposure of these dogs to infective third stage larvae in the sand/dirt exercise run/pens, which are ideal for hookworm transmission (Ridley et al., 1994). The combination of large numbers of animals and high transmission rates produces large effective populations of worms and increases the probabilities that resistance mutations will occur (Gilleard, 2006; Redman et al., 2015). Racing greyhounds are also treated extremely frequently with multiple different anthelmintics (e.g., fenbendazole, ivermectin, pyrantel) throughout their lives (Ridley et al., 1994). The intervals between these treatments often are less than the pre-patent period for hookworms, which will minimize the amount of refugia. Thus, genetically-resistant worms surviving treatment will have a large reproductive advantage, and the lack of refugia will lead to a rapid increase in their frequency (Martin et al., 1981; van Wyk, 2001). This combination of factors is known to place heavy selection pressure for drug resistance in nematodes (Wolstenholme et al., 2004), and is very similar to the epidemiological factors that have led to high levels of MDR in nematodes of sheep and goats, worldwide (Kaplan and Vidyashankar, 2012).

Given the fact that resistance is likely to have developed independently to each drug class, and the fact that MDR *A. caninum* to all three major anthelmintic classes seem to be virtually ubiquitous in racing greyhounds in the USA, one must ask the question; “why has resistance to any single drug class not been reported previously in the USA greyhound population”? To answer this question, we must examine how veterinarians typically manage hookworm infections in dogs. Typically, when a dog presents to a veterinarian with a fecal positive for hookworms, the dog is treated with one or more drugs from the benzimidazole, macrocyclic lactone or tetrahydropyrimidine classes. If the dog then tests positive again in a future exam, the infection is attributed to reinfection or reactivation of encysted/arrested larvae (larval leak). Consequently, the same treatment regimen is often repeated, or the veterinarian may choose to use a drug from a different drug class. One thing that was never done by small animal clinicians in the past, is measuring the efficacy of the treatment in a fecal egg count reduction test (FECRT) (Jimenez Castro and Kaplan, 2020; Kaplan, 2020), by performing both pre- and post-treatment FEC. As a result, anthelmintic resistance (AR) is not diagnosed, and most often is not even considered as a likely cause of the recurrent hookworm infections. Therefore, as resistance evolves and leads to more recurrent hookworm infections, veterinarians typically treat more often, and rotate and/or combine drugs. But they do not perform FECRT to measure the efficacy of the various drugs administered. Thus, as long as one drug remains efficacious, the problem will appear to be managed, and recurrent infections will continue to be attributed to reinfection or reactivation of encysted/arrested larvae. However, once MDR to all drugs evolves, it is no longer possible to manage the infections, and the problem of anthelmintic resistance becomes more obvious. Our data demonstrate evidence of very high IC_50_ and IC_95_ values for benzimidazoles and the macrocyclic lactones. With regards to pyrantel, no *in vitro* or molecular assays currently exist for measuring resistance. However, in every suspected MDR case we have treated with pyrantel pamoate there is virtually no efficacy based on FEC reduction, (Jimenez Castro et al., 2019), and in a controlled efficacy study using Worthy 4.1F3P, pyrantel pamoate yielded an efficacy of only 23% (Jimenez Castro et al., 2020).

Regarding benzimidazole resistance, the Sanford, FL kennel applied the greatest benzimidazole selection pressure of any of the kennels, treating all dogs twice a month with albendazole, and all dogs tested had F167Y (TTC>TAC) frequencies of at least 70%. When comparing the phenotypic and genotypic data for benzimidazole, the IC_95_ yielded a significant correlation (*p* = 0.04) but the IC_50_ did not (*p* = 0.08). The lack of significance for the IC_50_ may be due to low power, as a consequence of only having 15 samples with both types of data. However, this finding is consistent with our previous work where we found that the IC_95_ was more appropriate for discriminating susceptible vs resistant isolates using the EHA (Jimenez Castro et al., 2019). Interestingly, there were two samples, Dl 13 and Dl 14 that had IC_95_ values 13 and 18 times higher than the susceptible isolate from our previous work (Jimenez Castro et al., 2019), but had a resistant F167Y SNP allele frequency of only 5%, and 0%, respectively. This lack of correlation between phenotype and genotype raises three possible explanations: (1) the EHA has a high interassay variability, (2) there are mutations, other than at codons 167, 198 and 200, that are involved with resistance to benzimidazole drugs, or (3) there are loci other than β -tubulin that are involved with resistance to benzimidazole drugs. However, in on our previous work we tested multiple isolates and biological replicates and had rather low interassay variability (Jimenez Castro et al., 2019). Additionally, we previously measured a >100-fold increase in the EHA IC_50_ in the Worthy isolate following treatment with fenbendazole, but the SNP allele frequency remained relatively unchanged (Jimenez Castro et al., 2019). Together, these findings lend support to the hypothesis that there are non-β-tubulin mutations that are involved in resistance to benzimidazole drugs. Evidence of this has already been reported in *Caenorhabditis* spp. where a quantitative trait loci that did not overlap with β -tubulin genes was identified in two genetically divergent isolates (Zamanian et al., 2018). Also, using genome wide association mappings in *C. elegans*, novel genomic regions independent of *ben-1* and other β-tubulin loci were correlated with resistance to albendazole (Hahnel et al., 2018). Additionally, disparity in responses in *C. elegans* to fenbendazole and albendazole showed evidence that the former could have additional targets beyond β-tubulin, such as genes that encode β-tubulin interacting proteins (Dilks et al., 2020). Thus, our observations demand further study.

Our previous work demonstrated that the LDA provided excellent discrimination between susceptible and resistant isolates for the MLs (Jimenez Castro et al., 2019). In the current study the median IC_50_ value was >83 times higher than the susceptible isolate value from our previous work. This is both extremely high and an underestimation, since the IC_50_ of many samples could not be accurately measured given that they exceeded the highest concentration tested. Furthermore, of all the samples that had LDA assays with IC_50_ values >1000 nM, all but one had moxidectin as the last treatment administered, and that one group was administered moxidectin in the previous treatment. Macrocyclic lactones, particularly, ivermectin, have been used intensively by the greyhound industry for parasite control for the past several decades (Ridley et al., 1994). However, to our knowledge, moxidectin, which is a substantially more potent member of this drug class (Prichard et al., 2012), has only recently started to be used on greyhound farms and kennels. In *H. contortus*, ivermectin resistant worms that are naïve to moxidectin are typically killed at very high efficacy following administration of moxidectin (Craig et al., 1992; Oosthuizen and Erasmus, 1993); however, once moxidectin is used regularly in an ivermectin-resistant population, resistance to moxidectin can develop rapidly (Kaplan et al., 2007). Evidence suggests this same pattern is occurring in hookworms as well.

When examining the FEC data for moxidectin and albendazole by timeframe post-treatment, we found significant differences for both drugs, but in a different pattern. For the group of samples collected following albendazole treatment, the 10-21 (*p* = 0.0003) and >21 days (*p* = 0.0004) categories had a statistically significant higher mean FEC when compared to the <10 days category. This finding is consistent with our previous observations in the Worthy isolate, where we documented a temporary suppression on worm fecundity following treatment with fenbendazole (Jimenez Castro et al., 2019; Jimenez Castro et al., 2020). In contrast, for the dogs treated with moxidectin, there was no difference between the <10 and 10-21 day timeframes, but the >21day period had significantly higher FEC (*p* = 0.0002) and (*p* < 0.0001), respectively. This is also consistent with our previous observations where no reduction in fecundity was seen in moxidectin-resistant *A. caninum* following treatment with ML drugs. The higher EPG in the >21day period is likely due to reinfection and/or reactivation of arrested larvae.

The almost ubiquitous presence of MDR worms in recently retired greyhounds, combined with the demise of the greyhound racing industry and increasing numbers of greyhound adoptions, poses a serious risk to the health of pet dogs. From 2009-2019, the number of dog parks increased by 74% in the USA (TPL, 2019), and a recent survey showed that the prevalence of *A. caninum* in dogs visiting these parks (Stafford et al., 2020) was more than 70% higher as compared to the prevalence in all pet dogs recorded over the same general timeframe (Sweet et al., 2021). These relative prevalence data should not be surprising; a fecal pile deposited by a 30 kg dog with an *A. caninum* FEC of ∼1000 EPG will contain approximately 500,000 eggs. If not picked up, tens to hundreds of thousands of infective larvae are likely to contaminate the surrounding soil from this one defecation. Consequently, there is a high probability of transmission to other dogs visiting the dog park, and once infected, usual anthelmintic treatment of these dogs will have little efficacy. The end result will be a continual cycle of infection and transmission that is not interrupted by usual monthly treatments with heartworm preventive products, or with anthelmintics administered specifically to treat the hookworm infections. When this is considered in light of the fact that resistance in *A. caninum* was not reported in greyhounds until the worms were already MDR, it seems highly likely that anthelmintic-resistant *A. caninum* are already quite common in pet dogs.

Finally, given these new alarming data, it is urgent that studies be performed to determine the prevalence and geographic distribution of drug-resistant *A. caninum* in the general pet dog population. Additionally, studies investigating the haplotype diversity of the susceptible and resistant alleles in *A. caninum* isolates from both greyhounds and the general pet dog population from different geographic regions are likely to provide deeper insights into the molecular epidemiology and the origin(s) of these MDR worms.

## Acknowledgements

We thank the staff at the participating greyhound kennels, adoption groups and veterinarians for their efforts and assistance, which made this project possible.

