## Supplementary figures and images for "Multiple drug resistance in hookworms infecting greyhound dogs in the USA"

### Supplementary figure 1

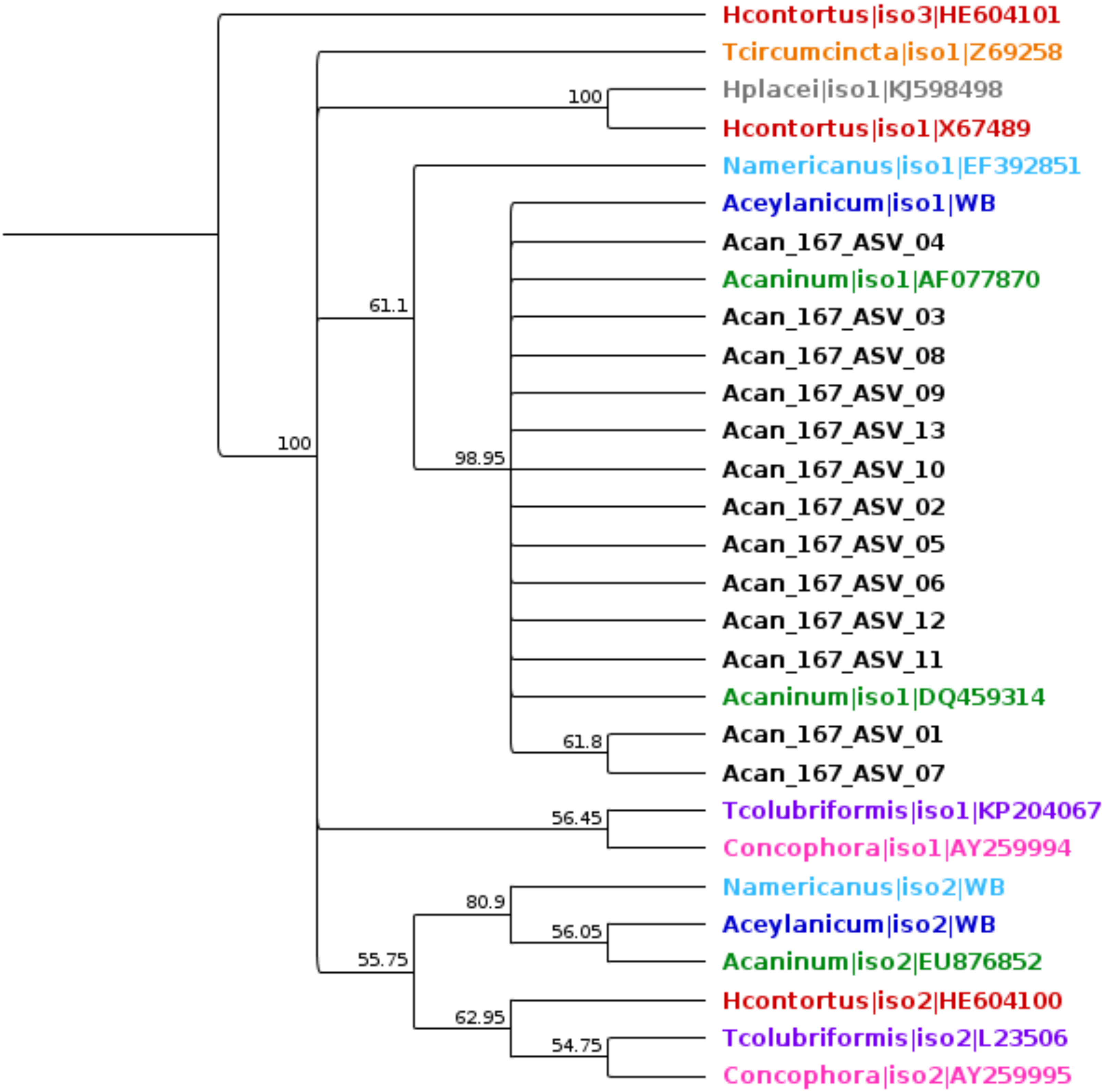

### Supplementary Figure 2

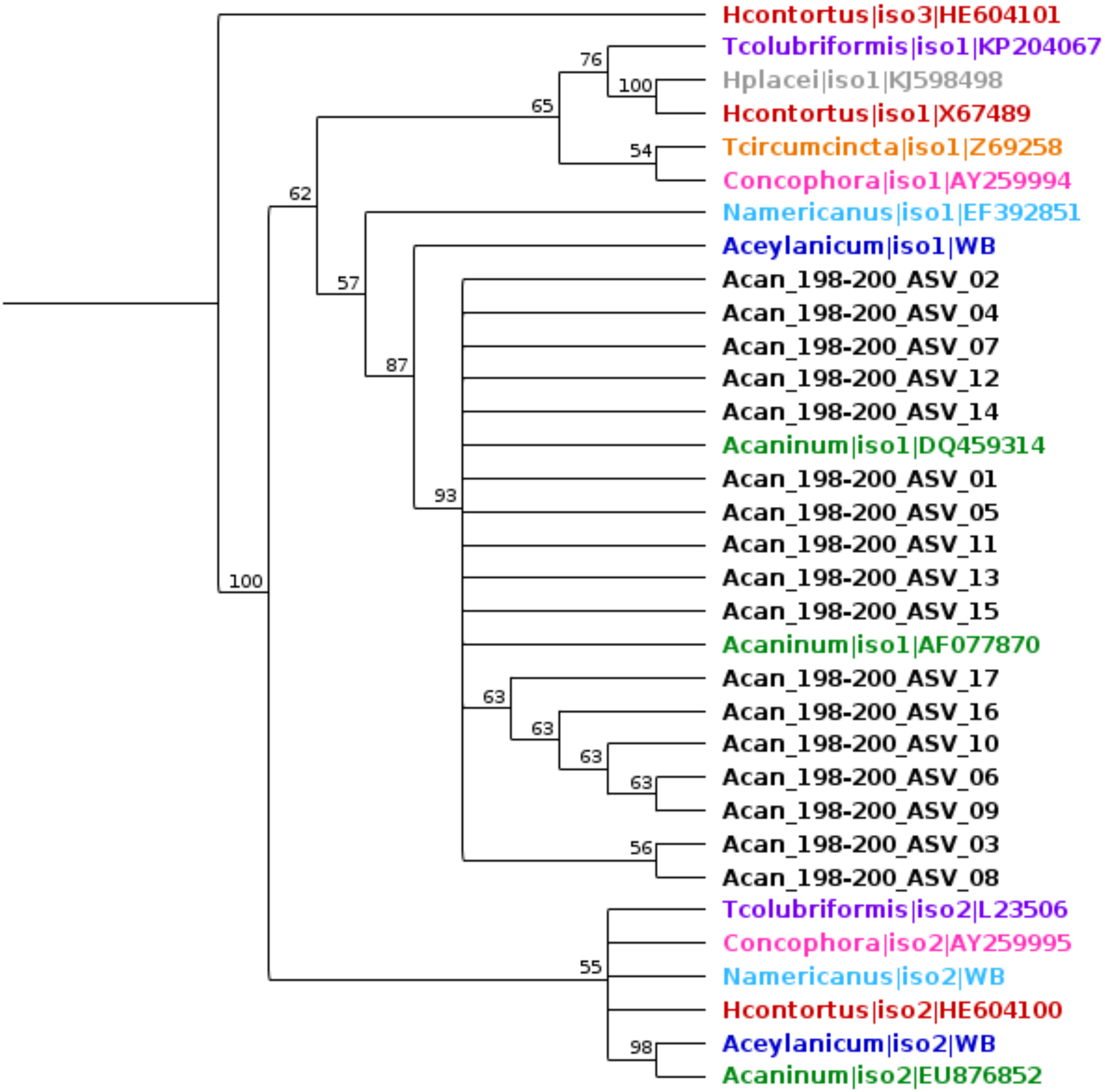
